# NRDE2 Interacts with an Early Transcription Elongation Complex and Widely Impacts Gene Expression

**DOI:** 10.1101/2025.08.20.671253

**Authors:** Marina Srbic, Chaïmaa Belhaouari, Raoul Raffel, Laurine Lemaire, Jerome Barbier, Julie Bossuyt, Charbel Akkawi, Xavier Contreras, Rosemary Kiernan

## Abstract

NRDE2 is a highly conserved protein implicated in post-transcriptional gene silencing in *Schizosaccharomyces pombe* and *Caenorhabditis elegans* and has been shown to modulate splicing in mammals. To explore whether NRDE2 participates in additional processes in human cells, we performed tandem affinity purification followed by proteomic analysis of NRDE2 from nuclear extracts of HEK293T and HeLa cells. Our analysis confirmed the interaction of NRDE2 with its well-characterized partner, the MTR4 helicase (MTREX), as well as with multiple splicing factors. Notably, we also identified interactions with chromatin-associated proteins involved in transcription, including the Polymerase-Associated Factor 1 (PAF1) complex and elongating forms of RNA polymerase II (RNAPII). To further investigate NRDE2 function, we conducted RNA-seq following its transient depletion. Differential expression analysis revealed that loss of NRDE2 alters the expression of thousands of genes. Consistent with earlier reports, we observed splicing defects, particularly intron retention; however, our results indicate that the impact of NRDE2 on intron retention is more extensive than previously recognized. Moreover, intron retention was frequently associated with reduced mRNA expression. Together, these findings suggest that NRDE2 associates with both the transcriptional and splicing machineries and plays a broader role in RNA processing than previously appreciated.

## Introduction

NRDE2 (nuclear RNAi-defective 2) is a conserved protein found in eukaryotes ranging from fission yeast to humans but absent in budding yeast. It was first identified in *Caenorhabditis elegans*, where it participates in nuclear RNA interference (RNAi), a process that mediates gene silencing through small RNAs, including siRNAs (small interfering RNAs) ^1^. In *C. elegans*, the NRDE pathway maintains heritable RNAi-mediated silencing. Nuclear-localized siRNAs guide NRDE factors to nascent pre-messenger RNAs (pre-mRNAs), leading to H3K9me3 deposition and inhibition of RNA polymerase II (RNAPII) during transcription elongation^1–3^. Together with NRDE-3, NRDE2 is required for the recruitment of NRDE-1 to pre-mRNAs and chromatin, thereby mediating small RNA–dependent silencing through H3K9 methylation ^4^. Notably, the conserved MTR4 helicase was shown to interact with NRDE2 and contribute to NRDE2-dependent co-transcriptional gene silencing ^5^.

In fission yeast, the NRDE2 homolog Nrl1 forms a complex with Mtl1 (the mammalian homolog MTREX, hereafter referred to as MTR4) and Ctr1, and interacts with core splicing factors ^6 7^. The Nrl1–Ctr1–Mtl1 complex has been proposed to regulate the splicing of cryptic introns and to target unspliced transcripts for degradation by the nuclear exosome. Nrl1 directs transcripts with cryptic introns to promote heterochromatin formation at developmental genes and retrotransposons ^6^.

In mammals, NRDE2 has been implicated in diverse cellular processes, including pre-mRNA splicing and the maintenance of genome integrity. Proteomic studies have identified numerous interactions with splicing factors, and loss of NRDE2 results in splicing defects, particularly retention of weakly spliced introns, in mouse and human cells ^8,9^. Loss of NRDE2 also leads to DNA damage and R-loop accumulation ^10^. More recent studies suggest a role for NRDE2 in DNA damage responses, potentially through interactions with repair pathways that safeguard gene expression under stress conditions ^11^. As in *S. pombe* and *C. elegans*, the principal binding partner of mammalian NRDE2 is the MTR4 helicase ^5,6,10,12^. Importantly, NRDE2 interaction with MTR4 is essential for NRDE2 protein stability ^5^. NRDE2 has been reported to inhibit exosome activity by restraining MTR4 function ^12^. Specifically, NRDE2 binding locks MTR4 in a closed conformation, preventing its interaction with both the exosome and cofactors, such as the cap-binding complex (CBC) and ZFC3H1 ^12^. Interestingly, the NRDE2-MTR4 interaction is required for self-renewal in mouse embryonic stem cells ^12^.

NRDE2 has also been implicated in human disease. It was identified as a biomarker in circulating tumor cells of patients with metastatic melanoma, based on genome-wide analysis of copy-number aberrations and loss-of-heterozygosity ^13^. More recently, NRDE2 was reported as a novel susceptibility gene for hepatocellular carcinoma (HCC) ^11^. Mechanistically, NRDE2 binds to casein kinase 2 (CK2) subunits and promotes holoenzyme assembly and activity. This NRDE2-dependent enhancement of CK2 activity facilitates MDC1 phosphorylation and DNA damage responses. A rare-variant, NRDE2-p.N377I, largely abolished this function and sensitized HCC cells to poly(ADP-ribose) polymerase (PARP) inhibitors ^11^. Additionally, NRDE2 has been shown to contribute to infection by Kaposi’s Sarcoma-Associated Herpesvirus (KSHV). In this context, NRDE2 prevents degradation of late viral transcripts by inhibiting RNA decay pathways ^14^.

Taken together, these findings suggest that NRDE2 plays an evolutionarily conserved role in splicing regulation. However, its precise molecular functions, particularly in mammalian systems, remain incompletely understood. Here, we identify novel NRDE2 interactors, including not only splicing factors but also previously unrecognized chromatin-associated partners linked to early elongating RNAPII complexes. Transcriptome analyses revealed that loss of NRDE2 alters expression of numerous genes, with clear evidence of splicing defects, especially intron retention. Nonetheless, many changes in gene expression appear to occur independently of splicing. These observations suggest that NRDE2 may exert additional roles in regulating gene expression beyond splicing.

## Results

### NRDE2 interacts with transcription-associated proteins and RNA processing factors

To investigate NRDE2 function, we generated Hela S3 and HEK293T cell lines stably expressing NRDE2 with tandem Flag and HA tags on the N-terminus (F.H-NRDE2). In HeLa S3 cells, F.H-NRDE2 was expressed following transduction with a lentiviral vector, as described previously ^15^ whereas in HEK293T cells the tag was inserted at the endogenous *NRDE2* locus using CRISPR-Cas9 technology ^16^. We first confirmed expression of F.H-NRDE2 in both cell lines and verified that a known NRDE2 partner, MTR4, was present in F.HNRDE2 immunoprecipitates (Supplementary Fig. 1a, b). Nuclear extracts of F.H-NRDE2-HeLa S3 or F.HNRDE2-HEK293T were immunoprecipitated using anti-Flag antibody and the precipitates were immunoblotted using anti-HA and anti-MTR4 antibodies (Supplementary Fig. 1a, b). These results demonstrated that F.H-NRDE2 was properly expressed and retained interaction with a known partner in both HeLa S3 and HEK293T cells.

To identify NRDE2 interactants in each cell type, nuclear extracts were subjected to tandem affinity purification using sequential anti-Flag followed by anti-HA antibodies, as described previously ^15,16^. Eluates were analyzed by tandem mass spectrometry (Taplin facility, Harvard University, Boston) and interactants ≥2 peptides and fold change (FC) >2 compared to control HeLa S3 or HEK293T cells were selected. In HeLa S3 cells Flag.HA-NRDE2 specifically interacted with 102 proteins, while 81 interactants were detected in HEK293T (Supplementary Table 1). Among these, 20 proteins were detected in both cell lines (Figure 1a and Supplementary Table 2). Gene ontology (GO) analysis revealed that NRDE2 interacted with proteins involved in RNA processing and splicing in both HeLa S3 (Figure 1b) and HEK293T cells (Figure 1c). Notably, a higher proportion of chromatin-associated interactants were detected in HeLa S3 cells compared to HEK293T. GO network analysis further indicated that NRDE2 interactants are predominantly associated with RNA processing, particularly RNA splicing (Supplementary Figure 2).

**Figure 1.**
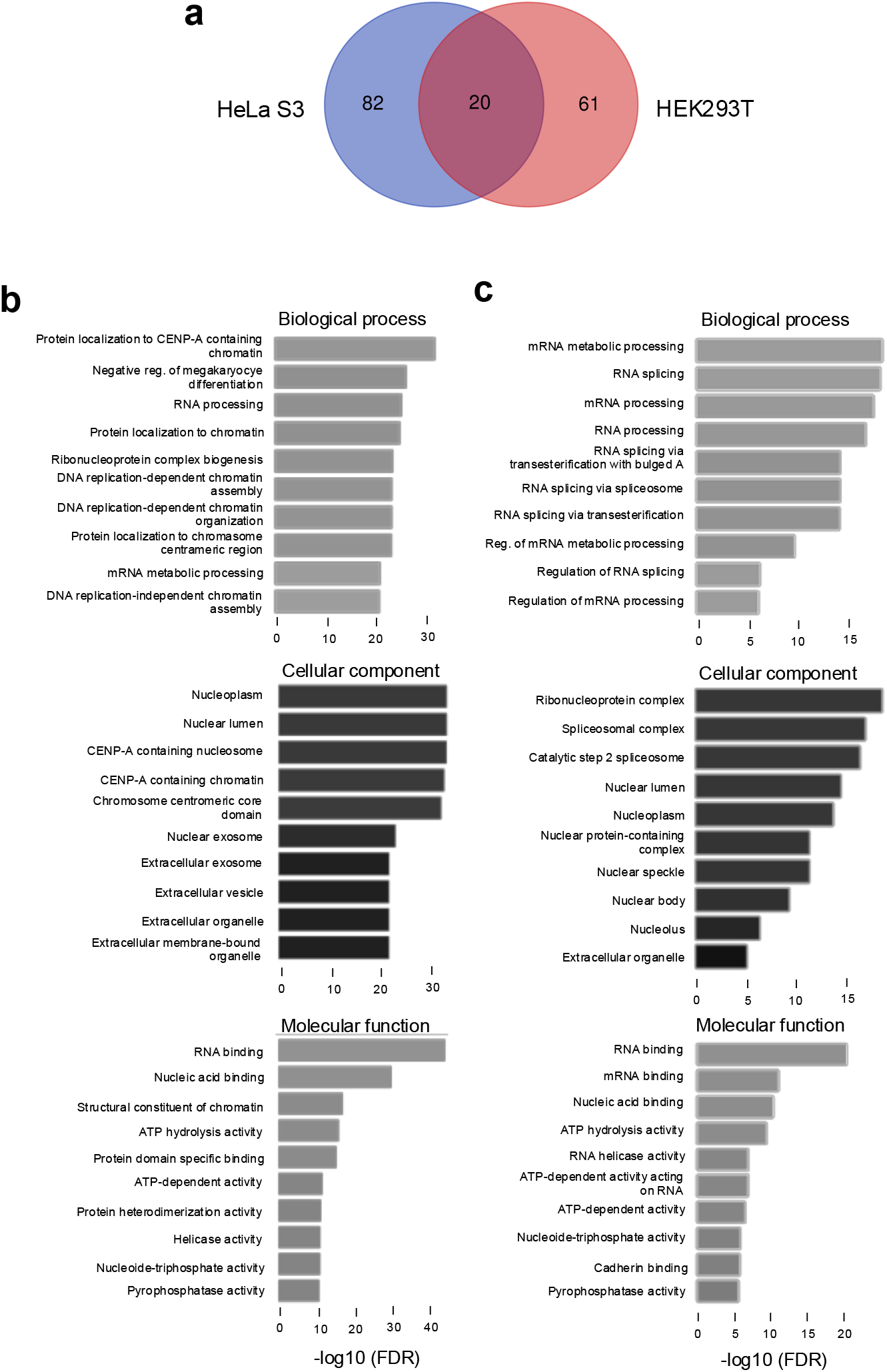
NRDE2 interacts with proteins found in the nucleus and chromatin-associated proteins that are implicated in RNA processing. a. Venn diagram showing the overlap between NRDE2 interactants detected in FH-NRDE2-expressing HeLa S3 or FH-NRDE2-expressing HEK293T cells, as indicated. b. GO analysis of NRDE2 interactants detected in FHNRDE2-expressing HeLa S3. c. GO analysis of NRDE2 interactants detected in FH-NRDE2-expressing HEK293T.

We next tested whether the F.H-NRDE2 interactants identified by proteomics could be detected in association with endogenous, untagged NRDE2. Several interactants representing distinct nuclear protein complexes were analyzed by co-immunoprecipitation using nuclear extracts from parental HEK293T cells. As expected, interaction with MTR4, the most abundant NRDE2 interactant identified in both cell types, was confirmed (Figure 2a). Interaction with additional RNA helicases, including DDX15 and DDX17, were also validated. Although splicing factors were represented by relatively few peptides in the NRDE2 interactome, interactions with EFTUD2, SFPQ, and SNRNP200 were confirmed. Likewise, interactions with RNA-binding proteins such as SAF-B, ILF2, and ILF3 were validated. Notably, Topoisomerase I (TOP1), a DNA gyrase, was identified as an NRDE2 interactor and its association with endogenous NRDE2 was also confirmed (Figure 2a).

**Figure 2.**
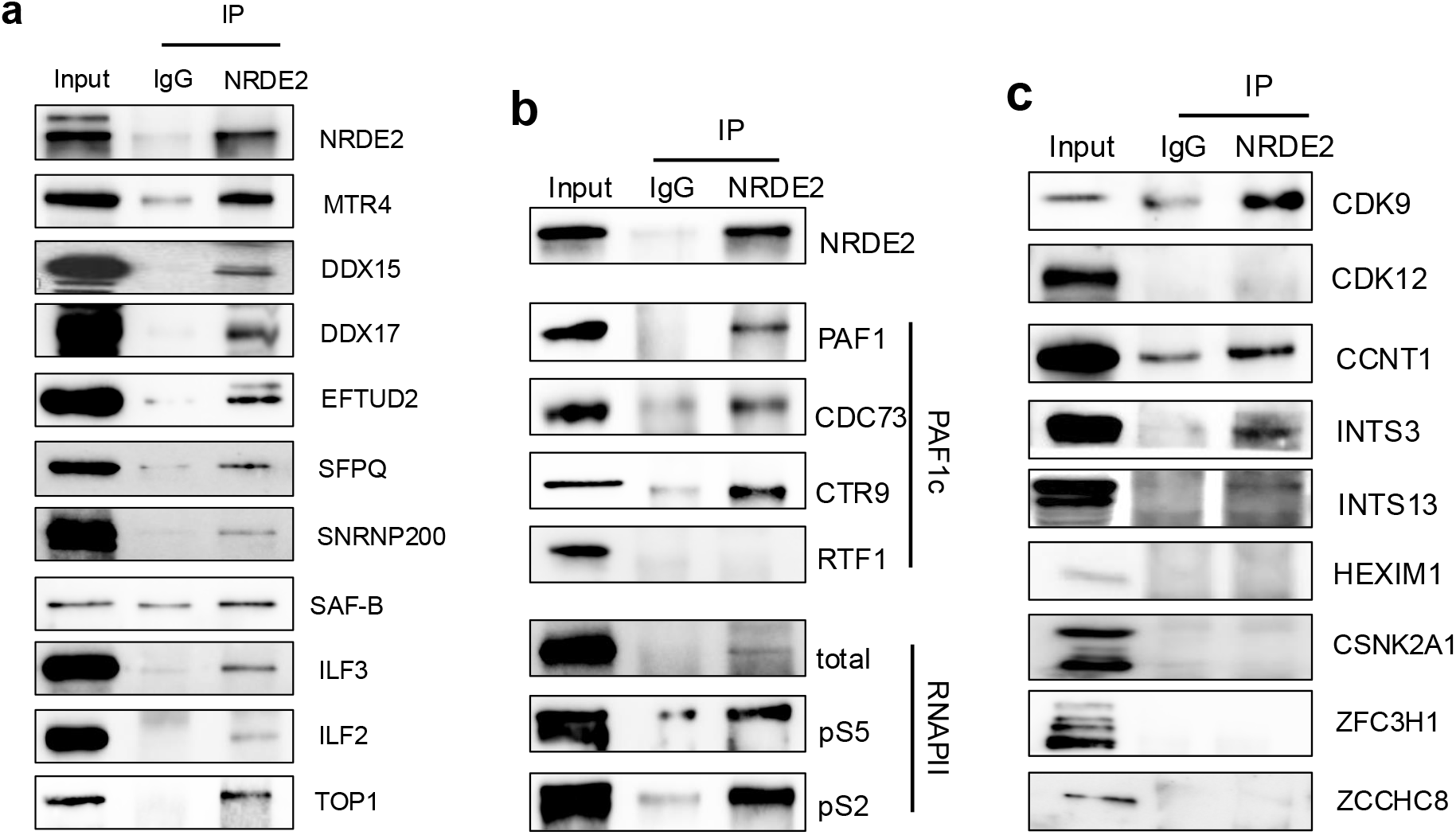
NRDE2 interacts with proteins implicated in transcription and mRNA processing. a. Co-immunoprecipitation analysis of endogenous NRDE2 and interactants identified by mass spectrometry in either HeLa S3, HEK293T or both cell types. HEK293T nuclear extracts were immunoprecipitated using an IgG control or NRDE2 antibodies, as indicated. Immunoprecipitates and an aliquot of nuclear extract (input, 5%) were analyzed by SDS-PAGE followed by immunoblot using the antibodies indicated on the figure. b. Co-immunoprecipitation analysis of endogenous NRDE2 and subunits of PAF1c identified by mass spectrometry, as well as with different forms of RNAPII. HEK293T nuclear extracts were immunoprecipitated using an IgG control or NRDE2 antibodies, as indicated. Immunoprecipitates and an aliquot of nuclear extract (input, 5%) were analyzed by SDS-PAGE followed by immunoblot using the antibodies indicated on the figure. c. Co-immunoprecipitation analysis of endogenous NRDE2 and RNAPII-associated proteins implicated in transcription. HEK293T nuclear extracts were immunoprecipitated using an IgG control or NRDE2 antibodies, as indicated. Immunoprecipitates and an aliquot of nuclear extract (input, 5%) were analyzed by SDS-PAGE followed by immunoblot using the antibodies indicated on the figure.

We also noted that both HeLa S3 and HEK293T FH-NRDE2 interactomes contained subunits of the multi-subunit Polymerase-Associated Factor 1 complex (PAF1c) although the specific subunits identified differed between the two cell types. The HEK293T interactome contained the CTR9 subunit, whereas the HeLa S3 interactome included PAF1, CDC73, and WDR61 (Supplementary Table 2). To test whether NRDE2 interacts with endogenous PAF1c, we performed co-immunoprecipitation assays. NRDE2 was found to associate with PAF1, CDC73, and CTR9, but not with the highly labile RTF1 subunit (Figure 2b), suggesting that NRDE2 primarily interacts with the core PAF1 complex.

Because PAF1 directly associates with transcribing RNA polymerase II (RNAPII) ^17,18^, we next asked whether NRDE2 could also be detected in complex with RNAPII. NRDE2 interacted modestly with total RNAPII, which includes both active and inactive forms. Strikingly, NRDE2 displayed stronger interactions with transcriptionally engaged RNAPII phosphorylated on serine 5 of the C-terminal domain (RNAPII pS5), which marks initiation and early transcription, as well as with RNAPII phosphorylated on serine 2 (RNAPII pS2), which marks elongation (Figure 2B). These findings indicate that NRDE2 associates with actively transcribing RNAPII, as well as with the PAF complex, which is known to co-ordinate histone modifications required for transcriptional activity.

Since NRDE2 interacted with actively transcribing forms of RNAPII, we next asked whether it might also associate with additional factors involved in transcription elongation. Endogenous NRDE2 was found to interact with CDK9 and cyclin T1, subunits of the positive transcription elongation factor b (P-TEFb) complex, which acts during the early stages of elongation (Figure 2c). In contrast, NRDE2 did not interact with CDK12, which regulates elongation across gene bodies (Figure 2c). P-TEFb is normally sequestered in an inactive complex by 7SK RNA and HEXIM1 ^19^. Interestingly, although NRDE2 interacted with CDK9 and cyclin T1, no interaction was detected with HEXIM1 (Figure 2c), suggesting that NRDE2 preferentially associates with the active form of P-TEFb.

We also tested whether NRDE2 interacts with subunits of the Integrator complex, which has been implicated in RNAPII pausing near transcription start sites ^20–22^. Indeed, NRDE2 was found to interact with INTS3 and INTS13 subunits (Figure 2c), suggesting that NRDE2 associates with the holo-Integrator complex. Together, these results demonstrate that NRDE2 interacts with multiple factors involved in active RNAPII transcription, particularly during early elongation.

NRDE2 has also been reported to associate with casein kinase 2 (CK2) subunits and to promote assembly of the CK2 holoenzyme ^11^. Consistent with this, CSNK2A1 and CSNK2A2 were identified in the F.HNRDE2 interactome from HeLa S3 cells. However, we were unable to confirm these interactions by co-immunoprecipitation in HEK293T cells (Figure 2c). This discrepancy may reflect differences between mass spectrometry–based purification and co-immunoprecipitation methods.

In addition, neither PAXT nor NEXT subunits were detected in NRDE2 interactomes from HeLa S3 or HEK293T cells. Co-immunoprecipitation likewise failed to detect interactions between NRDE2 and ZFC3H1 (PAXT subunit) or ZCCHC8 (NEXT subunit) (Figure 2c). These results are consistent with previous findings that NRDE2 prevents MTR4 helicase from associating with exosome-linked complexes ^12^. Nevertheless, several nuclear exosome subunits were identified as NRDE2 interactors in HeLa S3 cells (Supplementary Tables 1 and 2). Of note, a robust interaction between NRDE2 and EXOSC10 has been reported previously ^9^.

### NRDE2 affects expression and mRNA splicing at a subset of genes

Because NRDE2 interacts with transcription-associated proteins, and has been reported to modulate mRNA processing, including splicing ^8,9^, we next assessed its effect on the transcriptome. NRDE2 expression was reduced by RNAi (Figure 3a). Transcriptome analysis revealed that loss of NRDE2 had a broad impact on gene expression: approximately 13,000 genes were differentially expressed following NRDE2 depletion (Figure 3b). Among these, 6,239 genes were downregulated, while 6,439 genes were upregulated (FC > 1.5, *p* < 0.05). Representative genome browser tracks of downregulated genes (*SLC1A4* and *ARPIN*) and upregulated genes (*PLEKHB1*and *BTN3A2*) are shown in Figure 3c. These results demonstrate that NRDE2 exerts a substantial influence on the transcriptome.

**Figure 3.**
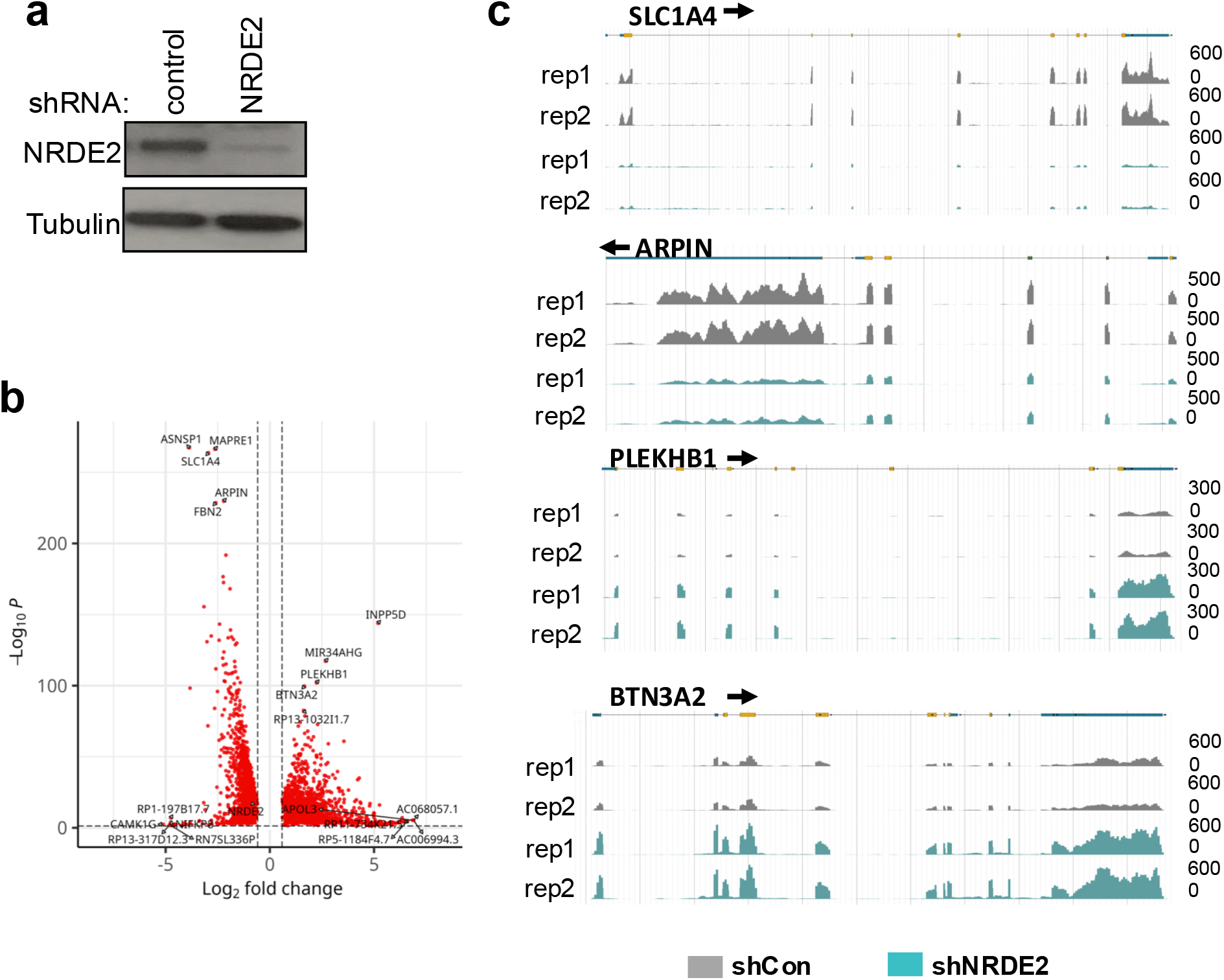
NRDE2 modulates the expression of a subset of genes. a. HeLa cells transduced with lentiviral particles expressing shRNA against NRDE2 or a nontargeting control were harvested 5 days post-transduction. Extracts were analyzed by immunoblot using the indicated antibodies. b. Volcano plot showing the differential expression of mRNAs in extracts of HeLa cells transduced with lentiviral particles expressing shRNA against NRDE2 or a non-targeting control. c. Browser shots of RNA-seq signal from control or NRDE2 knock-down samples, as indicated, over representative genes. A schematic representation of the gene is shown above.

### NRDE2 regulates intron retention and additional splicing events

NRDE2 has previously been shown to influence the splicing of short, GC-rich introns with relatively weak 5’ and 3’ splice sites in both mouse and human cells ^8,9^. To assess whether similar splicing defects could be detected in our system, we analyzed RNA-seq data from control and NRDE2 knockdown cells. Because intron retention (IR) is a known target of NRDE2, we first examined IR using IRFinder ^23^. Consistent with earlier studies, NRDE2 was found to suppress IR at a subset of genes ^8,9^. Notably, we detected substantially more NRDE2-regulated IR events in HeLa cells compared with MDA-MB-231 cells ^9^. In HeLa cells, 2298 deregulated IR events were detected using IRFinder (default parameters) (Figure 4a) whereas only 178 were reported previously in MDA-MB-231 cells ^9^. Despite this difference, the relative proportion of events was similar, representing 9% of all IR events in HeLa cells and 8% in MDA-MB-231 cells. Among deregulated IR events in HeLa cells, 94% were suppressed by NRDE2, consistent with previous observations ^9^.

**Figure 4.**
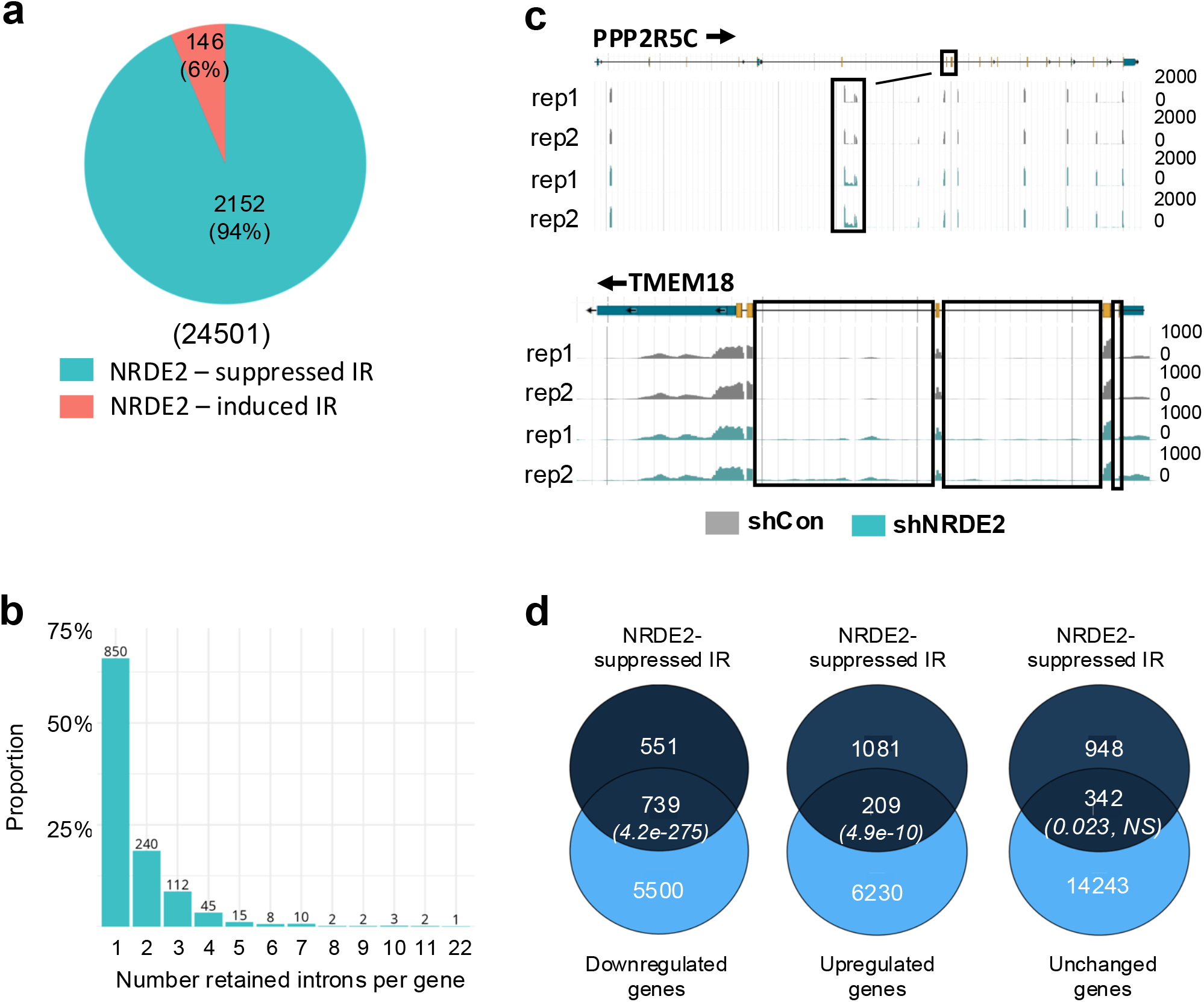
NRDE2 affects splicing at a subset of genes. a. Piechart showing the number of differentially retained introns (NRDE2-suppressed or NRDE2-induced) detected by IRFinder in RNA-seq samples from control compared to NRDE2 knock-down cells. b. Distribution analysis of NRDE2-suppressed IR showing the number of retained introns per gene. Numbers above the bars show the number of NRDE2-suppressed IR events. c. Browser shots of RNA-seq signal from control or NRDE2 knock-down samples, as indicated, over representative genes. A schematic representation of the gene is shown above. Boxed regions show NRDE2-suppressed IR. d. Venn diagrams showing the overlap between genes exhibiting NRDE2-suppressed IR and down-regulation (left), upregulation (middle) or unchanged (right) expression. P-values of the overlap (Fisher’s exact test) are shown in brackets on the Venn diagrams.

Analysis of the distribution of NRDE2-suppressed IR events revealed that most genes (66%) displayed retention of only a single intron following NRDE2 depletion (Figure 4b). Retention of up to four introns per gene accounted for 86% of all NRDE2-suppressed IR events. Thus, loss of NRDE2 led to retention of a limited number of introns per gene and was restricted to a specific subset of genes. Examples include *PPP2R5C*, which displayed retention of a single intron, and *TMEM18*, which showed retention of multiple introns (Figure 4c).

To determine whether NRDE2 also regulates other splicing events beyond intron retention, we analyzed RNA-seq data using Suppa2 ^24^. This analysis revealed that NRDE2 contributes to additional splicing outcomes, including skipped exons, mutually exclusive exons, alternative last exons, alternative first exons, and alternative 5’ and 3’ splice site usage (Supplementary Figure 3). While each of these events was deregulated upon NRDE2 depletion, they were less frequent than IR. Specifically, deregulated events represented approximately 5–7% of the total number of each event in control cells, compared with 12% for IR (Supplementary Figure 3).

We next examined the relationship between NRDE2-suppressed IR and differential gene expression (Figure 4d). Comparison of RNA-seq differential expression analysis with NRDE2-suppressed IR revealed a strong association with reduced mRNA levels (Figure 4d, left Venn diagram, *p=4.2e-275*, Fisher’s exact test). Notably, 57% of NRDE2-suppressed IR occurred in down-regulated genes. A smaller but significant proportion (16%) was associated with upregulated genes (Figure 4d, middle Venn diagram, *p=4.9e-10*, Fisher’s exact test), while 27% of events were found in genes with unchanged expression (Figure 4d, right Venn diagram, *p=0.023*, Fisher’s exact test). Thus, NRDE2-suppressed IR is frequently linked to altered gene expression, particularly downregulation.

Importantly, however, expression of nearly 12,000 genes was altered following NRDE2 depletion in a manner apparently independent of intron retention. This suggests that NRDE2 may exert broader regulatory effects on gene expression beyond its role in controlling IR.

## Discussion

NRDE2 is a highly conserved protein implicated in splicing regulation from fission yeast to humans, and in small RNA-dependent gene silencing in lower organisms. To better characterize human NRDE2, we characterized its interactome in two human cell types, HeLa S3 and HEK293T. Consistent with previous studies ^5,10,12^, the major interactant of NRDE2 in both systems was the RNA helicase, MTR4 (MTREX). While many interactants were shared between the two cell types, others were cell type–specific. Gene ontology (GO) analysis confirmed strong enrichment for proteins involved in RNA processing and splicing, consistent with prior reports showing that NRDE2 interacts with numerous splicing factors ^8,9^. In addition, interactants identified in HeLa S3 were enriched for chromatin-associated factors, which may reflect differences in chromatin extraction efficiency compared to HEK293T. Notably, several subunits of the RNAPII-associated PAF1 complex were identified in both systems, and interactions with PAF1, CDC73, and CTR9 were validated by coimmunoprecipitation (Figure 2b). In line with previous reports that RTF1 is not a stable component of the human PAF1 complex ^25–27^, it was not detected in association with NRDE2.

Our analyses further revealed that NRDE2 interacts with RNAPII, particularly with the phosphorylated forms involved in transcription initiation (pS5) and elongation (pS2) (Figure 2b). NRDE2 also associated with CDK9 and cyclin T1, subunits of the positive transcription elongation factor P-TEFb, but not with CDK12 or HEXIM1 (Figure 2c). Since P-TEFb controls the transition from promoter-proximal pausing to productive elongation, whereas CDK12 acts later during elongation and termination [28], these interactions suggest that NRDE2 may act primarily at an early step of transcription elongation near promoter regions. Supporting this, NRDE2 was also found to interact with subunits of the Integrator complex (INTS3 and INTS13) that regulate RNAPII pausing and termination in promoter-proximal regions [20–22].

Loss of NRDE2 had widespread consequences for gene expression. Of the 27,263 genes analyzed, nearly half (12,678 genes, 46.5%) were differentially expressed upon NRDE2 depletion (Figure 3). Roughly equal numbers of genes were up- or downregulated. In agreement with previous reports [8,9], NRDE2 suppressed intron retention (IR) in a subset of pre-mRNAs. In HeLa cells, NRDE2 modulated almost 10% of all detected IR events (Figure 4a), a higher fraction than reported previously in MDA-MB-231 cells [9]. Importantly, NRDE2-suppressed IR was significantly associated with changes in gene expression, most frequently with transcript downregulation (Figure 4d). Nevertheless, thousands of genes displayed altered expression without evidence of IR changes, suggesting that NRDE2 also influences gene expression through splicing-independent mechanisms.

Taken together, our results confirm that NRDE2 interacts with RNA processing and splicing factors and uncover novel associations with factors involved in early transcription elongation. Our RNA-seq analysis not only reinforces the role of NRDE2 in suppressing intron retention but also reveals that this effect is more widespread in human cells than previously appreciated. Furthermore, the finding that nearly half of all expressed genes are affected by NRDE2 depletion underscores its broad impact on gene expression. While splicing defects likely contribute to this regulation, most differential expression appears to occur independently of splicing, raising the possibility that NRDE2 might modulate transcription through its interactions with elongating RNAPII and associated complexes. Future work will be required to dissect the mechanisms by which NRDE2 coordinates transcription and RNA processing to ensure proper gene expression.

## Materials and Methods

### Cell Culture and Reagents

HeLa and HeLa S3 cells were grown in Dulbecco modified Eagle’s minimal essential medium (DMEM) (Sigma-Aldrich D6429), supplemented with 10% FCS (Eurobio Scientific, CVFSVF00-01), and containing 1% Penicillin streptomycin (Sigma P4333). HEK-293T cells (ATCC, CRL11268) were grown in Hepes-modified DMEM (Sigma-Aldrich, D6171), supplemented with 10% FCS (Eurobio Scientific, CVFSVF00-01), 2 mM GlutaMAX (Gibco, 35050-061) and containing 1% penicillin-streptomycin (Sigma-Aldrich, P4333). Both cell lines were grown in a humidified incubator at 37 °C with 5% CO2.

### Antibodies

Antibodies used in this study are shown in Table S3.

### Generation of FH-NRDE2-expressing cells

FH-NRDE2-expressing HeLa S3 cells were generated by transduction of lentiviral vectors expressing FHNRDE2, as previously described ^15^. Briefly, plasmid encoding pOZ-Flag-HA-NRDE2 (pOZ-FH-NRDE2) was cloned using pOZ-N-FH vector. Lentiviral particles expressing FH-NRDE2 were produced in HEK293T cells by transfecting plasmids using calcium-phosphate. HeLa S3 were transduced using Polybrene infection/transfection reagent (Sigma-Aldrich^®^, TR-1003), according to the manufacturer’s instructions. After 7 days, selection of transduced cells was carried out by magnetic affinity sorting with antibody against IL2 to achieve a pure population. HeLa S3 stably expressing NRDE2 (FH-NRDE2 HeLa S3) were confirmed by Western blot using anti-HA and anti-Flag antibodies.

FH-NRDE2-expressing HEK293T cells were generated by CRISPR-Cas9 mediated editing of endogenous NRDE2 gene. An sgRNA targeting the NRDE2 gene around the ATG translation start site was cloned in pSpCas9 (BB)−2A-GFP plasmid (Addgene #48138). The plasmid was then transfected into HEK-293T cells along with a single-stranded oligodeoxynucleotide (ssODN) (Table S4) harboring the Flag-HA sequence flanked by homology sequences to NRDE2 around the cleavage site. Single cells were isolated and amplified. HEK293T clones expressing Flag-HA NRDE2 were identified by PCR and confirmed by sequencing as well as Western blot using anti-HA and anti-Flag antibodies.

### RNAi

Production of short hairpin RNA (shRNA)-expressing lentiviral particles was performed as described previously ^28^ using plasmids expressing shRNAs targeting NRDE2 (Sigma-Aldrich MISSION shRNA, TRCN0000422527) or a nontargeting control (Addgene, plasmid #1864). For knockdown experiments, HeLa cells were transduced with lentiviral particles and harvested 5 days later.

### NRDE2 Protein Complex Purification

NRDE2 protein complex was extracted from nuclear extracts of HeLa S3 or HEK-293T cells stably expressing Flag-HA-NRDE2. Briefly, cells were seeded in 150 mm culture dishes the day prior to protein extraction. Cytoplasmic proteins were extracted using a mild lysis buffer [10 mM Hepes pH 7.9, 10 mM KCl, 0.1 mM EDTA pH 8.0, 2 mM MgCl2, 1 mM DTT, protease and phosphatase inhibitors (Halt™, Thermofisher 78438)]. The cell pellet was incubated for 10 min on ice, adding 0.07% NP-40 and incubating for an additional 15 min on ice. After centrifugation (2000 x g, 5 min, 4 °C), the cytoplasmic fraction was collected and discarded. The pellet was washed with ice-cold PBS supplemented with protease and phosphatase inhibitors. After centrifugation (2000 x g, 5 min 4 °C), the supernatant was discarded leaving the packed nuclear volume (PNV). For extraction of nuclear soluble proteins, nuclei were resuspended drop-wise in 1x PNV of hypotonic buffer [20 mM Hepes pH 7.9, 20 mM NaCl, 1 mM EDTA pH 8.0, 1.5 mM MgCl2, 10% glycerol, 1 mM DTT, protease and phosphatase inhibitor (Halt™, Thermofisher 78438)], followed by the addition of 1x PNV of high salt buffer (20 mM Hepes pH 7.9, 800 mM NaCl, 1 mM EDTA pH 8.0, 1.5 mM MgCl2, 10% glycerol, 1 mM DTT, protease and phosphatase inhibitor (Halt™, Thermofisher 78438)]. Samples were rotated on a wheel for 20 min at 4 °C, followed by centrifugation (20 min 16,000 x g, 4 °C). The supernatant containing the nuclear extracts was collected.

FH-NRDE2 complex was purified by two-step affinity chromatography, as described previously (dx.doi.org/10.17504/protocols.io.kgrctv6). Sequential Flag and HA immunoprecipitations were performed on equal amounts of proteins. Silver-staining was performed according to the manufacturer’s instruction (Silverquest, Invitrogen). Mass spectrometry was performed at Taplin facility, Harvard University, Boston, MA.

### Coimmunoprecipitation analysis

Coimmunoprecipitation was performed using nuclear extracts of HEK-293T cells as described previously ^29^. Briefly, cells were washed twice with PBS and then resuspended in Cytoplasmic Lysis Buffer (CLB) [10 mM Tris, pH = 7.4; 10 mM NaCl; 3 mM MgCl_2_; 0.5% NP-40, complemented with protease and phosphatase inhibitors (Halt™, Thermofisher 78438)]. Nuclei were immediately pelleted by centrifugation at 300 × *g* for 4 min at 4°C. The supernatant (cytoplasmic fraction) was discarded. Another CLB wash was performed before lysing the nuclei in a Nuclear Lysis Buffer (NLB) [20 mM Tris (pH = 8), 3 mM KCl, 150 mM NaCl, 1 mM MgCl_2_ and 0.2% Triton™ X-100, protease and phosphatase inhibitors (Halt™, Thermofisher 78438)]. After a 30 min incubation on ice, extracts were cleared by centrifugation at 14 000 rpm for 15 min at 4°C and the supernatant containing nuclear protein extracts was collected. For immunoprecipitation, nuclear protein extracts (250 μg) were incubated with antibodies (2 μg) overnight on a rotating wheel at 4°C. Protein A (Invitrogen, Dynabeads™ Protein G, 10009D) conjugated beads were added and the mixtures were incubated for 30 min at room temperature on a rotating wheel. Following 3 washes with NLB, Samples were resuspended in protein sample loading buffer (1x Laemmli buffer), incubated for 5 min at 95°C and analyzed by Western blotting using the antibodies shown in Table S2.

### RNA-seq and statistical analysis

For RNA-seq, total RNA was extracted from HeLa cells using TRIzol (Thermo Fisher Scientific) according to the manufacturer’s instructions. Paired end sequencing of polyA+ selected mRNA (20M clean reads per sample, 125 bp reads) was carried out by BGI Genomic Services (BGISEQ-500, NGS platform) in triplicates. Statistical analyses were performed as follows:

Quality Control and Pre-processing of Sequencing Data: Initial quality control of raw sequencing data was performed using FastQC (v0.11.9), and results were summarized using MultiQC (v1.11.dev0). Adapter trimming and quality filtering were conducted using fastp (v0.23.2) with default parameters. Reads were also screened against contamination sources using FastQ Screen (v0.14.1) to ensure sample integrity.

Read Alignment and Reference Genome: Processed reads were aligned to the human reference genome GRCh38 (Ensembl release 84) using STAR aligner (v2.7.10a) with a two-pass alignment strategy. The genome was indexed using STAR with the parameters --sjdbOverhang 50.

Post-alignment Processing: Duplicate reads arising from PCR amplification were removed using SAMtools rmdup (v1.12). Aligned BAM files were sorted by genomic coordinates and indexed using SAMtools index (v1.12).

Quantification of Gene Expression: Gene-level quantification was performed using featureCounts (v2.0.3) with the Ensembl GRCh38 release 84 GTF annotation file, counting reads mapping uniquely to exons.

Differential Expression Analysis: Differential gene expression analysis was carried out with DESeq2 (v1.30.1) in R/Bioconductor. Data normalization and differential expression were assessed using negative binomial generalized linear models, and results were visualized with PCA plots, MA plots, volcano plots, and heatmaps for quality control and biological interpretation.

Intron Retention Analysis: Intron retention was quantified using IRFinder (v2.0.0) ^23^. Differential intron retention between experimental conditions was assessed using DESeq2 integrated within IRFinder, identifying statistically significant intron retention events. Additional differential splicing events were assessed using Suppa2 ^24^.

Additional Bioinformatics Tools and Analyses: Additional genomic manipulations and analyses were performed using bedtools (v2.29.1), and genomic visualization was facilitated using deepTools (v3.5.1). Workflow automation and reproducibility were managed using Snakemake (v7.32.4). Gene ontology was performed using ShinyGO (0.82) (https://bioinformatics.sdstate.edu/go/). Enrichments in Venn diagrams were performed using Fisher’s exact test.

## Supporting information

support figs and tables

## Supplementary Materials

The following supporting information can be downloaded at: www.mdpi.com/xxx/s1, Figure S1: FH-NRDE2 interacts with MTR4; Figure S2: NRDE2 interactants are implicated in RNA processing and chromatin-associated processes; Figure S3: NRDE2 modulates splicing. Table S1: NRDE2 interactants identified by mass spectrometry in HEK293T or HeLa S3 stably expressing Flag-HA-NRDE2.; Table S2: NRDE2 interactants identified in both HEK293T or HeLa S3 Flag-HA-NRDE2expressing cells or in HEK293Tor HeLa S3-Flag-HA-NRDE2-expressing cells specifically. Table S3: Antibodies used in this study. Table S4: Oligonucleotides used in this study.

## Author Contributions

Conceptualization, Marina Srbic, Xavier Contreras and Rosemary Kiernan; Data curation, Marina Srbic, Raoul Raffel and Rosemary Kiernan; Formal analysis, Marina Srbic, Chaimaa Belhaouari, Raoul Raffel, Laurine Lemaire and Rosemary Kiernan; Funding acquisition, Rosemary Kiernan; Investigation, Marina Srbic, Chaimaa Belhaouari, Laurine Lemaire, Jérôme Barbier, Julie Bossuyt and Charbel Akkawi; Methodology, Chaimaa Belhaouari, Raoul Raffel, Laurine Lemaire and Jérôme Barbier; Project administration, Julie Bossuyt, Charbel Akkawi and Rosemary Kiernan; Software, Raoul Raffel; Supervision, Xavier Contreras and Rosemary Kiernan; Validation, Marina Srbic, Chaimaa Belhaouari, Laurine Lemaire, Jérôme Barbier, Julie Bossuyt and Charbel Akkawi; Writing – original draft, Rosemary Kiernan; Writing – review & editing, Xavier Contreras and Rosemary Kiernan.

## Data Availability Statement

The data supporting the findings of this study are available from the corresponding authors upon request. All raw sequencing data and processed files (aligned reads, count tables, and analysis outputs) have been deposited in GEO under accession number GSE300903.

## Acknowledgments

The authors thanks members of the Gene Regulation laboratory for helpful discussions.

