## Supplementary material for "NRDE2 Interacts with an Early Transcription Elongation Complex and Widely Impacts Gene Expression": support figs and tables

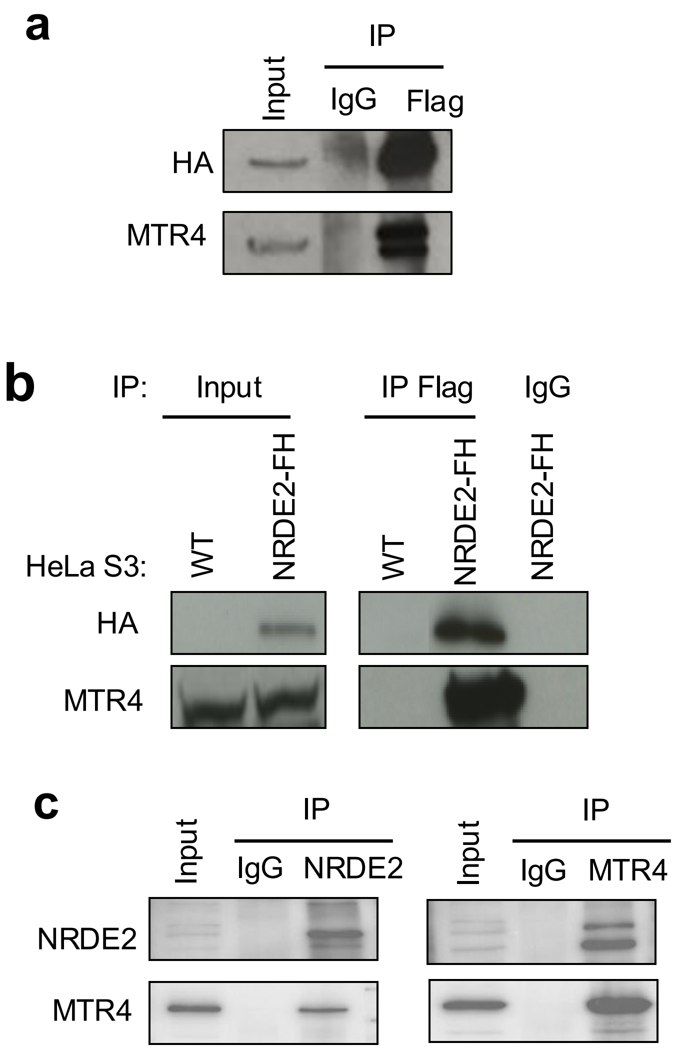

**Supplementary Figure 1.** FH-NRDE2 interacts with MTR4

**a.** Nuclear extracts from FH-NRDE2-HEK293T cells were used for immunoprecipitation using anti-IgG control or anti-Flag antibody, as indicated. Immunoprecipitates and an aliquot of extract (input, 5%) were analyzed by Western blot using anti-HA or anti-MTR4 antibodies, as indicated. **b.** Nuclear extracts from HeLa S3 or FH-NRDE2-HeLa S3 cells were used for immunoprecipitation using anti-IgG control or anti-Flag antibody, as indicated. Immunoprecipitates and an aliquot of extract (input, 5%) were analyzed by Western blot using the antibodies indicated. **c.** Nuclear extracts from FH-NRDE2-HEK293T cells were used for immunoprecipitation using anti-IgG control, anti-NRDE2 or anti-MTR4 antibody, as indicated. Immunoprecipitates and an aliquot of extract (input, 5%) were analyzed by Western blot using anti-anti-NRDE2 or anti-MTR4 antibodies, as indicated.

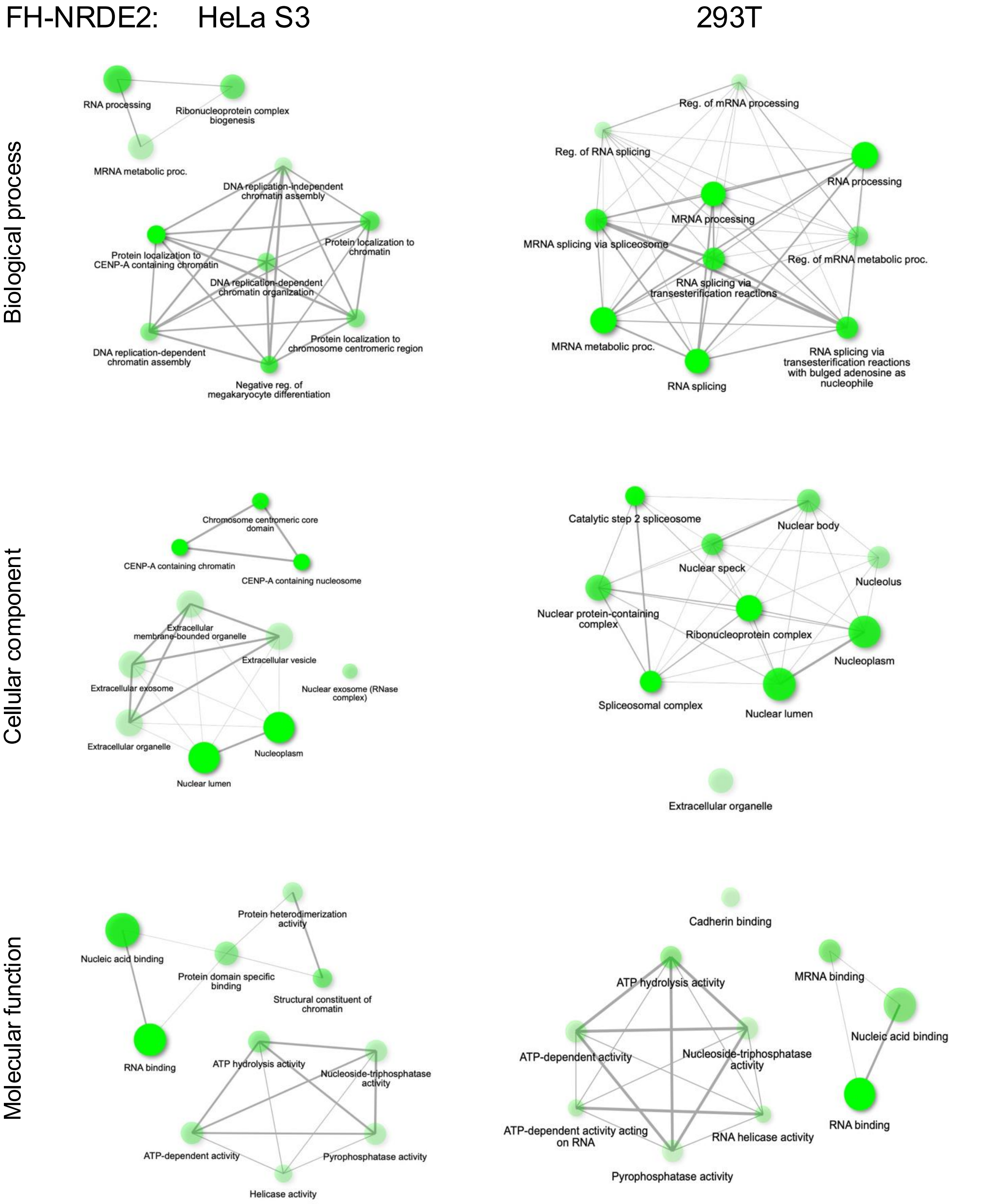

**Supplementary Figure 2.**

**Supplementary Figure 2.** NRDE2 interactants are implicated in RNA processing and chromatin-associated processes.

NRDE2 interactants identified from FH-NRDE2 HeLa S3 or FH-NRDE2 HEK293T cells, as indicated were analyzed using ShinyGO 0.82 (https://bioinformatics.sdstate.edu/go/) using default parameters. Gene ontology analyses of biological process, cellular components and molecular function of NRDE2 associated proteins are shown. Plots show the relationship between enriched GO pathways. Pathways, that are depicted as nodes, are shown as connected if they share 20% or more genes. Node colour intensity represents the significance of enriched gene sets. Node size represents the number of genes. Thickness of connecting lines represents the number of overlapping genes.

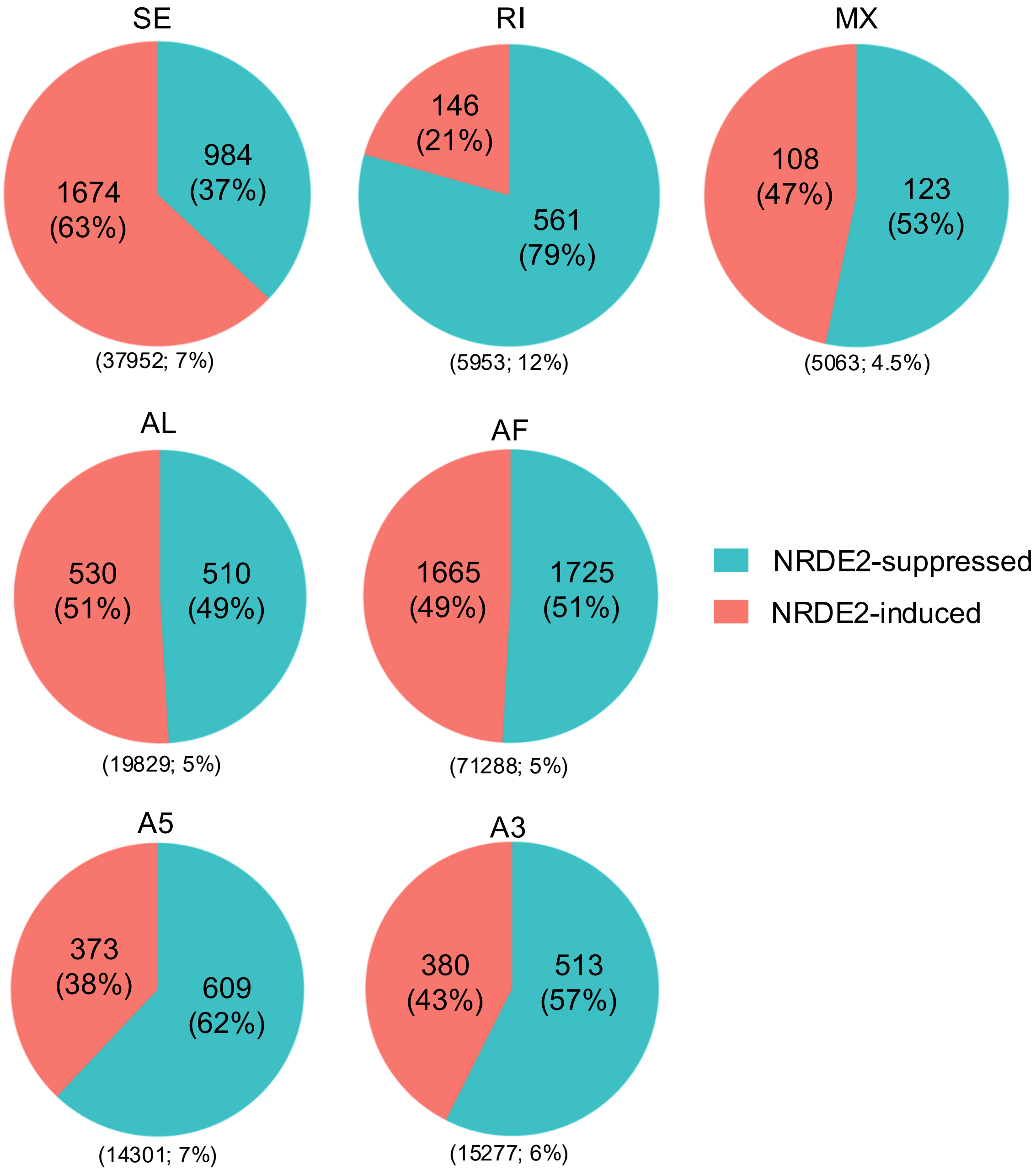

**Supplementary Figure 3.** NRDE2 modulates splicing.

**a.** Protein extracts from HeLa cells transduced with shRNA targeting NRDE2 or a non-targeting control were analyzed by Western blot using the indicated antibodies. **b.** Analysis of RNA-seq samples obtained from control or NRDE2 knock-down HeLa cells using Suppa2. Pie charts show the number of splicing events suppressed by NRDE2 (green) or induced by NRDE2 (red). Percentages in brackets show the proportion of the total number of differentially regulated events. The total number of splicing events detected in the samples and the percentage deregulated by NRDE2 knock-down are shown in brackets below the pie chart. Splicing events represented in the pie charts are skipped exons (SE), retained introns (RI), mutually exclusive exons (MX), alternative last exon (AL), alternative first exon (AF) alternative 5′ splice site (A5) and alternative 3′ splice site (A3).

**Supplementary Table 1**. NRDE2 interactants identified by mass spectrometry in HEK293T or HeLa S3 stably expressing Flag-HA-NRDE2.

| FH-NRDE2 HeLa S3 | | | | FH-NRDE2 HEK293T | | | |
|---|---|---|---|---|---|---|---|
| Protein ID | peptide count | Protein ID | peptide count | Protein ID | peptide count | Protein ID | peptide count |
| NRDE2 | 242 | H2BC21 | 3 | MTREX | 118 | NOVA1 | 2 |
| MTREX | 229 | HNRNPF | 3 | NRDE2 | 93 | NR3C1 | 2 |
| EXOSC10 | 67 | HNRNPH1 | 3 | PKM | 9 | NXN | 2 |
| HSPA8 | 37 | HNRNPK | 3 | HNRNPA3 | 8 | POLDIP3 | 2 |
| CSNK2A2 | 33 | HSPA9 | 3 | ACTB | 7 | PRAMEF20 | 2 |
| HSPA5 | 33 | NOP58 | 3 | ANXA2 | 7 | PRPF19 | 2 |
| CSNK2A1 | 31 | PRPF8 | 3 | NPM1 | 7 | PRSS33 | 2 |
| HNRNPM | 31 | SFN | 3 | HSPA8 | 6 | PTPRB | 2 |
| EXOSC2 | 25 | SNRPD2 | 3 | DBN1 | 4 | S100A14 | 2 |
| ACTA2 | 21 | PRPF8 | 3 | DDX17 | 4 | S100A16 | 2 |
| EXOSC7 | 18 | SFN | 3 | DDX5 | 4 | SAFB | 2 |
| MPHOSPH6 | 18 | SNRPD2 | 3 | DHX15 | 4 | SAFB2 | 2 |
| CDC73 | 16 | AKAP8 | 2 | EWSR1 | 4 | SERBP1 | 2 |
| EXOSC9 | 16 | AKAP8L | 2 | HNRNPR | 4 | SF3B1 | 2 |
| ACTB | 15 | CENPB | 2 | HSPA5 | 4 | SF3B2 | 2 |
| LMNA | 14 | CHAF1B | 2 | RBMX | 4 | SFPQ | 2 |
| PAF1 | 14 | CHD1 | 2 | AZGP1 | 3 | SNRNP200 | 2 |
| SERPINB4 | 13 | CUL9 | 2 | FABP9 | 3 | SNRNP40 | 2 |
| HSPA1A | 12 | DDX31 | 2 | HNRNPH1 | 3 | SRSF1 | 2 |
| ANXA1 | 11 | DDX3X | 2 | ILF2 | 3 | TAF15 | 2 |
| EXOSC8 | 10 | DDX52 | 2 | ILF3 | 3 | TERT | 2 |
| HSPA1L | 10 | DNAJA1 | 2 | POF1B | 3 | TOP1 | 2 |
| RUVBL2 | 10 | DNAJB6 | 2 | ANXA1 | 2 | TPM1 | 2 |
| TUBB | 10 | EXOSC5 | 2 | ASPRV1 | 2 | UBA52 | 2 |
| EXOSC3 | 8 | GTF3C4 | 2 | ATP1B3 | 2 | YBX1 | 2 |
| NPM1 | 8 | H2AZ2 | 2 | CEP112 | 2 | ZDHHC13 | 2 |
| ACTBL2 | 7 | HIST1H2AB | 2 | CEP152 | 2 | ZNF140 | 2 |
| ACTN1 | 7 | HNRNPD | 2 | CTR9 | 2 | ZNF248 | 2 |
| DDX5 | 7 | HSP90AB1 | 2 | DDX3X | 2 | | |
| EXOSC1 | 7 | HSP90B1 | 2 | DDX3Y | 2 | | |
| HIST1H4J | 7 | MAT2A | 2 | DDX47 | 2 | | |
| RUVBL1 | 7 | NCCRP1 | 2 | ECE2 | 2 | | |
| SNRNP200 | 7 | PCBP1 | 2 | EYA2 | 2 | | |
| TUBA1C | 7 | PCBP2 | 2 | FABP5 | 2 | | |
| EXOSC6 | 6 | PFKP | 2 | GGCT | 2 | | |
| HSPB1 | 6 | PFN1 | 2 | GTF2I | 2 | | |
| TGM1 | 6 | PRPF19 | 2 | HECTD1 | 2 | | |
| AHNAK | 5 | QSER1 | 2 | HNRNPAB | 2 | | |
| C1D | 5 | RBM39 | 2 | HNRNPC | 2 | | |
| DHX15 | 5 | RFC5 | 2 | HNRNPDL | 2 | | |
| RFC4 | 5 | SNRNP40 | 2 | HNRNPM | 2 | | |
| YWHAE | 5 | SNRPD1 | 2 | HRC | 2 | | |
| DDX17 | 4 | SNRPD3 | 2 | HSPA1A | 2 | | |
| GNB2L1/Rack1 | 4 | TBL3 | 2 | HSPA1L | 2 | | |
| HSP90AA1 | 4 | TCOF1 | 2 | HSPA2 | 2 | | |
| NOP56 | 4 | TPM3 | 2 | ISG15 | 2 | | |
| SERPINB3 | 4 | UBR5 | 2 | KIF14 | 2 | | |
| TYMP | 4 | VCP | 2 | KIF15 | 2 | | |
| UBC | 4 | WDR5 | 2 | LIMA1 | 2 | | |
| ATP5F1A | 3 | WDR61 | 2 | MPHOSPH6 | 2 | | |
| ATP5F1B | 3 | XRCC6 | 2 | MUL1 | 2 | | |
| CHD9 | 3 | YWHAZ | 2 | NAPRT | 2 | | |
| EXOSC4 | 3 | | | NCCRP1 | 2 | | |

**Supplementary Table 2**. NRDE2 interactants identified in both HEK293T or HeLa S3 Flag-HA-NRDE2-expressing cells or in HEK293T- or HeLa S3-Flag-HA-NRDE2-expressing cells specifically.

| FH-NRDE2 -expressing cell line | Number of proteins | Protein name |
|---|---|---|
| HEK293 and HeLa S3 | 20 | ACTB; ANXA1; DDX3X; DDX5; DHX15; DDX17; HNRNPH1; HNRNPM; HSPA1A; HSPA1L; HSPA5; HSPA8; MPHOSPH6; NCCRP1; NPM1; NRDE2; PRPF19; MTREX; SNRNP40; SNRNP200 |
| HEK293 only | 61 | ANXA2; ASPRV1; ATP1B3; AZGP1; CEP112; CEP152; CTR9; DBN1; DDX3Y; DDX47; ECE2; EWSR1; EYA2; FABP5; FABP9; GGCT; GTF2I; HECTD1; HNRNPA3; HNRNPAB; HNRNPC; HNRNPDL; HNRNPR; HRC; HSPA2; ILF2; ILF3; ISG15; KIF14; KIF15; LIMA1; MUL1; NAPRT; NOVA1; NR3C1; NXN, PKM; POF1B; POLDIP3; PRAMEF20; PRSS33; PTPRB; RBMX; S100A14; S100A16; SAFB; SAFB2; SERBP1; SF3B1; SF3B2; SFPQ; SRSF1; TAF15; TERT; TOP1; TPM1; UBA52, YBX1; ZDHHC13; ZNF140; ZNF248 |
| HeLa S3 only | 82 | ACTA2; ACTBL2; ACTN1; AHNAK; AKAP8; AKAP8L; ATP5F1A; ATP5F1B; C1D; CDC73; CENPB; CHAF1B; CHD1; CHD9; CSNK2A1; CSNK2A2; CUL9; DDX31; DDX52; DNAJA1; DNAJB6; EXOSC1; EXOSC2; EXOSC3; EXOSC4; EXOSC5; EXOSC6; EXOSC7; EXOSC8; EXOSC9; EXOSC10; GTF3C4; H2AZ2; H2BC21; HIST1H2AB; HIST1H4J; HNRNPD; HNRNPF; HNRNPK; HSP90AA1; HSP90AB1; HSP90B1; HSPA9; HSPB1; LMNA; MAT2A; NOP56; NOP58; PAF1; PCBP1; PCBP2; PFKP; PFN1; PRPF8; QSER1; RACK1; RBM39; RFC4; RFC5; RUVBL1; RUVBL2; SERPINB3; SERPINB4; SFN; SNRPD1; SNRPD2; SNRPD3; TBL3; TCOF1; TGM1; TPM3; TUBA1C; TUBB; TYMP; UBC; UBR5; VCP; WDR5; WDR61; XRCC6; YWHAE; YWHAZ |

**Table S3.** Antibodies used in this study.

| Antibody | Reference | Supplier | Dilution used (WB) |
|---|---|---|---|
| CCNT1 | A303-499A | Bethyl laboratories | 1/2000 |
| CDC73 | Nb200-184 | Novus Biological | 1/500 |
| CDK9 | sc-13130 | SCBT | 1/500 |
| CDK12 | 26816-1-AP | Proteintech | 1/2000 |
| CSNK2A1 | ab70774 | Abcam | 1/1000 |
| CTR9 | A301-395A | Bethyl laboratories | 1/2000 |
| DDX15 | sc-271686 | SCBT | 1/500 |
| DDX17 | sc-398168 | SCBT | 1/1000 |
| EFTUD2 | A300-957A | Bethyl laboratories | 1/2000 |
| HA | sc-7392 | SCBT | 1/1000 |
| HEXIM1 | A303-112A | Bethyl laboratories | 1/500 |
| Histone H3 | ab1791 | Abcam | 1/5000 |
| ILF2 | A303-147A | Bethyl laboratories | 1/2000 |
| ILF3 | A303-651A | Bethyl laboratories | 1/2000 |
| INTS3 | 16620-1-AP | Proteintech | 1/1000 |
| INTS13 | 19892-1-AP | Proteintech | 1/1000 |
| MTR4 | 12719-2-AP | Proteintech | 1/1000 |
| NRDE2 | PA5-100410 | Invitrogen | 1/500 |
| PAF1 | A300-173A | Bethyl laboratories | 1/1000 |
| RNAPII (pSer2) | A300-654A | Bethyl laboratories | 1/2500 |
| RNAPII (pSer5) | ab5408 | Abcam | 1/1000 |
| RNAPII (total) | sc-899 | SCBT | 1/500 |
| RTF1 | A300-179A | Bethyl laboratories | 1/10000 |
| SAF-B | sc-393403 | SCBT | 1/500 |
| SFPQ | sc-271796 | SCBT | 1/500 |
| SNRNP200 | sc-393170 | SCBT | 1/500 |
| SP1 | sc-59 | SCBT | 1/200 |
| TOP1 | A302-589A | Bethyl laboratories | 1/2000 |
| ZCCHC8 | A301-806A | Bethyl laboratories | 1/2000 |
| ZFC3H1 | A301-456A | Bethyl laboratories | 1/2000 |
| TUBULIN | T6199 | Sigma-Aldrich | 1/10000 |

**Table S4.** Oligonucleotides used in this study

| Target | Sequence 5’ – 3’ |
|---|---|
| **ssODN NRDE2** | CGCCGGCGCCATCCCTATAGAGAAGAACGGCGGTACGTCCTGTGGTCATGGACTACAAGGACGACGATGACAAGCTCGATGGAGGATACCCCTACGACGTGCCCGACTACGCCGGAGGACTCGAGGCGCTGTTCCCAGCCTTTGCGGGGCTTAGTGAGGCTCCCGATGGCGGGAG |
| **sgRNA** | Sense: 5'-CACCGGAGGTACGGCCTGTGGTCA-3’<br><br>Antisense: 5'-AAACTGACCACAGGCCGTACCTCC-3' |
